# A Probabilistic Approach To The Diagnosis Of Obstructive Sleep Apnea

**DOI:** 10.1101/080515

**Authors:** Raj Ramnani, Anshul Tripathi

## Abstract

In a novel approach to diagnose Obstructive Sleep Apnea, electronic components including an Arduino Mega, a Bluetooth Transceiver, an accelerometer, and a air-quality sensor, were put together to create a wearable that would detect the frequency of apnea events, diagnose the disorder, and sound an alarm when necessary. A primary consideration was to make the mechanism accessible and affordable, and in doing so, lower the number of cases of Obstructive Sleep Apnea that go undiagnosed due to the cost and inconvenience associated with the traditional diagnosis method—a Polysomnogram. Bluetooth capability was an additional consideration so that the device would transmit data directly to an android smartphone, eliminating the need for an additional output mechanism. The total cost of the device, quite surprisingly, did not exceed $30, and therein rendered the device an accessible, affordable mechanism for diagnosis. Tests of the device on diagnosed patients yielded data consistent with the diagnosis, with a few false positives as a result of the excessive sensitivity of the sensors.

## I. INTRODUCTION

Sleep apnea is a chronic sleep disorder which results in frequent pauses in breathing/shallow breathing during the sleep of a person. Obstructive sleep apnea is a type of sleep apnea caused when the throat muscles occasionally relax and constrict the airway, therefore preventing breathing. In other words, tissues in the upper airways approach and nearly come into contact with each other, blocking the inward flow of air. Symptoms of this sleep disorder can be general weakness during the daytime, headaches, snoring, etc.

The most common methods of diagnosis include polysomnogram [PSG] at sleep laboratories [overnight sleep studies] and sleep monitoring devices. Both of these courses of detection are very expensive and often inconvenient, and consequently, unobtainable by the financially weaker section of society.

The primary objective of this project was to create a device that has the ability to perform the following operations:

I. Diagnose a person with sleep apnea
II. Detect the number and frequency of apnea events during a sleep session, and consequently, can calculate the Apnea-Hypopnea Index [an indicator of the severity of the case of apnea in a given patient, defined as the apnea events per hour of sleep]
III. Trigger an alarm system to alert patients in case an apnea attack lasts too long.
IV. Yield data offering insight into the effects of Air Temperature, Humidity, and CO_2_ concentration on a patient’s apnea attacks.
V. Compatible with an Android phone, and therefore, have Bluetooth functionality

The aim of this project was to end up with an open source device that can be modified/enhanced and used for research purposes, as well as be used by a common man suffering from sleep apnea to diagnose the presence and severity of his condition. It is worth the noting that the device in no way aims to replace the use of a breathing aid, such as a Continuous Positive Airway Pressure [CPAP] device, used by patients already diagnosed with the disorder; rather we aim to make the diagnosis possible so that someone suffering from Sleep Apnea can seek immediate medical assistance.

An understanding of the mechanism of the pathophysiology of Obstructive Sleep Apnea was deemed integral to the creation of the device and is therefore detailed below:

As a result of inhalation, air enters the body through either the mouth or the nose and makes its way to the pharynx. The path thus far is termed the upper airway. Ordinarily, the back of the pharynx is not robust and would therefore collapse inward as a person inhales.

However, dilator muscles in place prevent this from happening and keep the airway open. Under abnormal circumstances entailing one or more of the following: Dysfunction of the upper airway dilator muscles, Disproportionate Pharyngeal Wall Compliance, Intrapharyngeal and Peripharyngeal Impediments. An Apnea event can occur, temporarily hindering breathing. Most events do not wake the person but some may cause them to intermittently gasp/snort for breath.

Obstructive Hypopnea can be defined as incomplete obstruction of the upper airway, which leads to shallow breathing. This leads to snoring indistinguishable from that not caused by apnea, rendering diagnosis laborious. The continuation of the blockage of the upper airway entails a positive feedback loop wherein the dearth of oxygen in the blood signals the diaphragm to contract and therein draw air into the lungs. As inhalation increases, the blockage is exacerbated and can lead to dire consequences unless the dilator muscles regain control.

## II. PARTS USED

The following parts have been used for the creation of this device:

Arduino Mega [$9.50]
HC05 Bluetooth Transceiver [$4.50]
MPU6050 [3-axis gyroscope + 3-axis accelerometer; $3]
5 Volt Piezo Buzzer [$0.15]
DHT11 [Temperature + Humidity sensor] [$1.50]
MQ-135 Air Quality Sensor [measures CO_2_ concentration; $4]
DS 1307 Real Time Clock [RTC] Module [$1.50] Capacitive Touch Sensor [$2.50]
SD Card Module [$2]
*All of these materials combined cost under $30.*

### Outline

The remainder of this paper is organized as follows. Section 3 contains a comprehensive description and analysis of the functions and workings of this contraption. Section 4 contains our test results.

## III FUNCTIONS AND WORKING

### Sleep Apnea Diagnosis/ Apnea Attack Detection

For the purpose of detection and diagnosis of sleep apnea, an accelerometer is used to measure chest movement. When the device is strapped around the chest, the accelerometer is in a position parallel to the chest plate.

The accelerometer measures the force acting on it in 3 directions; the x-axis [along one edge of the accelerometer], the y-axis [along the edge perpendicular to the x-axis edge], and the z-axis [perpendicular to the plane of the accelerometer]. When stationary and with its plane parallel to the surface of the earth [i.e. the person is lying flat on his back], the only force acting on the accelerometer is downward gravity [F_G_]. F_G_ is also along the z-axis and therefore, F_G__z_ = F_G_.

When a person inhales, his chest exerts an upward force [F_B__z_] perpendicular to the z-axis of the accelerometer. In the case that the person is not lying flat on his back, the accelerometer is at an incline. Since the accelerometer is always strapped to the chest, breathing always exerts a force along the accelerometer’s z-axis.

However, gravitational force vector acts downwards and is broken into components F_Gx_, F_Gy_, and F_Gz_. Therefore, in this case, the net gravitational force is calculated using the vector resolution formula:

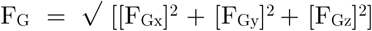

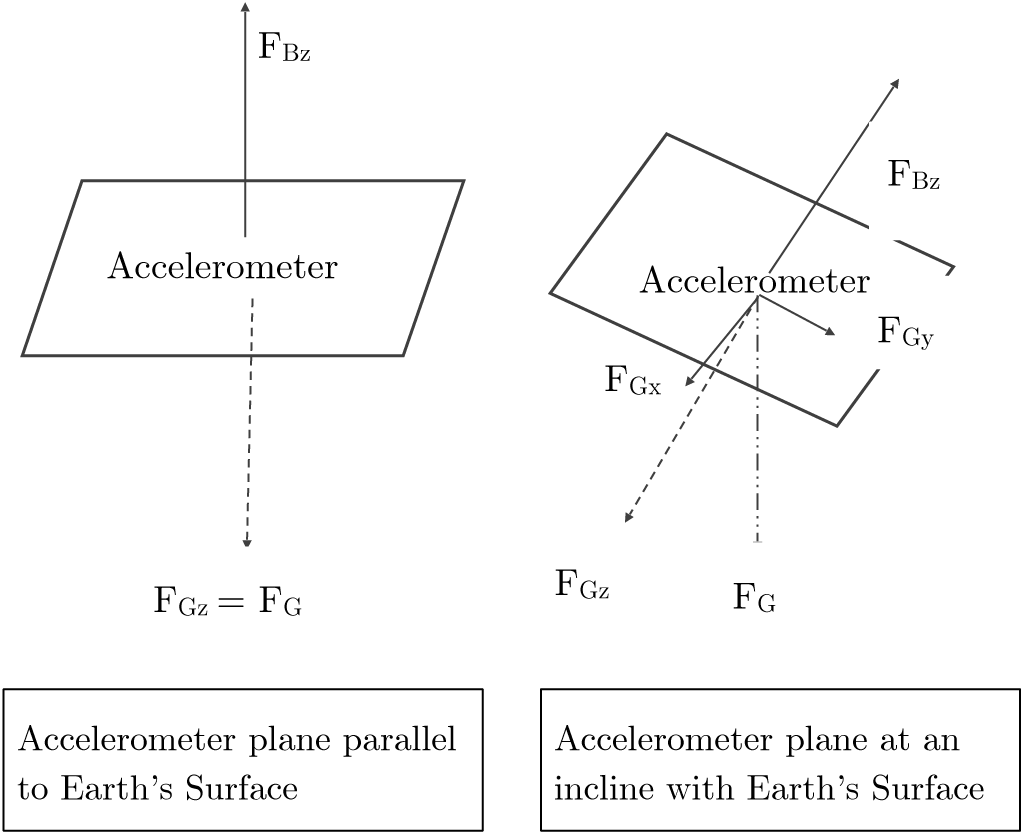

The algorithm that detects breathing is shown below:

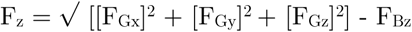

This algorithm entails that, when stationary, F_Bz_ = 0. However, during inhalation, when the chest exerts an upward force, F_B__z_ becomes greater than 0, and the value of F_z_ dips. This dip refers to one breath. For a person without sleep apnea, these dips are periodic.

However, if the accelerometer goes for a relatively longer period of time without the detection of such a dip, then it can be inferred that the person has stopped breathing and is suffering from an apnea attack. Such a period of invariance can be called an invariance event.

It is worth noting that hypopnea events [incomplete OSA, where the airway is only partially obstructed] will not trigger a dip in F_z_, so the device would be incapable of detecting and diagnosing hypopnea.

#### Self-Learning Algorithm

This algorithm is executed during the first run of the device and establishes the average frequency of inhalation. This algorithm allows for versatility by allowing it to be customized to different individuals with different breathing patterns.

#### Noise-Cancellation Algorithm

The accelerometer, when programmed, shows the values of force in terms of gravitational force along each axis. This means that when stationary and parallel to Earth, F_G__z_ ≈ 1.00 G, F_Gx_ ≈ 0.00 G, and F_Gy_ ≈ 0.00 G, where the numbers mean 1 G force, 0 G force, and 0 G force respectively.

However, because this value is only precise to 2 decimal places and can practically vary only between −1.00 G to 1.00 G [it is impossible for the chest to generate a force greater than gravity, i.e. 1.00 G], it is not very accurate

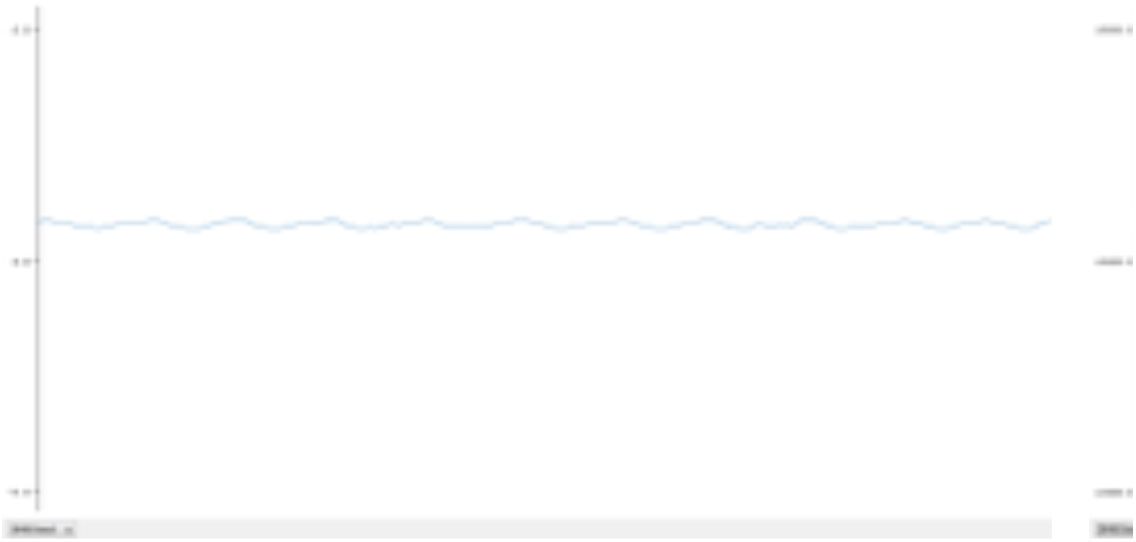

The dips in the wavy line above, which represent breathing, although visible, are very small in magnitude. Therefore, the preciseness needed to be increased. Registers are the memory locations where the accelerometer values are stored. Upon some research, we find out that the raw values being pulled from the registers are being divided by a scaling factor of 16384. Therefore, we can remove this division operation and receive values ranging from −16384 to +16384. Therefore, we obtain a precision that is just over 163 times greater than when the raw values are divided by the scaling factor.

However, owing to the variation and imperfection in the raw values, a great amount of noise is encountered.

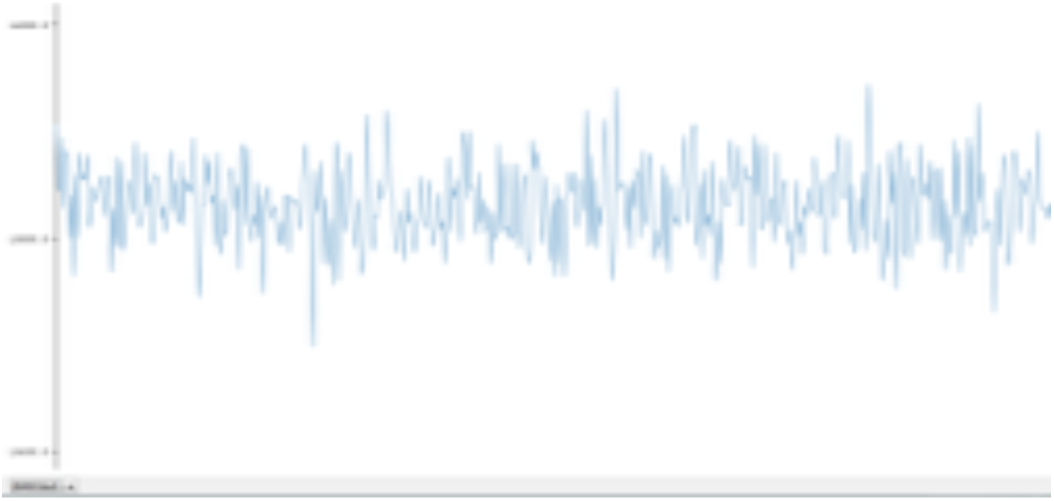

This noise is visible in the above graph. Such a great amount of noise makes pattern recognition and dip detection difficult. Therefore, to counteract this and smooth out the noise to a manageable level, a noise reduction algorithm is used. This algorithm is as follows:

**Let R_i_ be the i^th^ reading of the accelerometer. Let R_in_ be the new data point in the set of noise-reduced values**

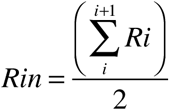

Using this simple noise reduction algorithm and recursively running it through the data points about 3, the noise is relatively lesser, and patterns more noticeable.

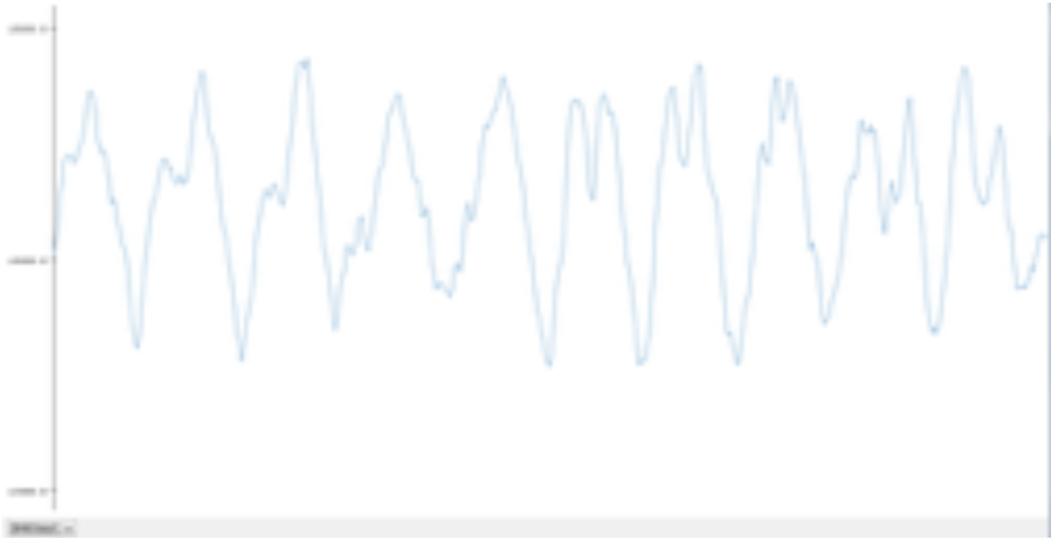

This is how pattern detection becomes easier and more precise.

### Calculation of Apnea-Hypopnea Index

The Apnea-Hypopnea index is used to quantify the severity of sleep apnea. It is defined as the total number of apnea events divided by the total number of hours of sleep.

In this device, this is accomplished with the help of a touch sensor and a real time clock [RTC] module. When the user touches the touch sensor once, an interrupt is triggered and the program starts running. A counter variable keeps track of the number of invariance events [a period of time during which dips in the graph are not detected]. Each invariance event points to an apnea attack.

After the user has finished sleeping and wakes up, he touches the sensor again. The total number of apnea events is then divided by the difference between the start time and end time. The result number gives us the AHI.

An AHI of >30 would signal excessive risk of acute myocardial infarction or a stroke and would alert the user accordingly.

A limitation of the method used is the failure to calculate a person’s Respiratory Disturbance Index [RDI]. The Respiratory Disturbance Index [RDI] indicates and any form of breathing irregularities and could possibly account for hypopnea-triggered deviations. As a result, one’s AHI is always equal to or lower than one’s RDI. Since the RDI isn’t calculated, risk posed by irregularities not in the form of described dip would go unidentified.

### Alarm Mechanism

If left unattended and allowed to continue for extended periods of time, apnea events have the potential to lead to a heart attack, a stroke, or even cardiac arrhythmia.

To prevent such a mishap from occurring, an alarm system has been incorporated into this device. If an invariance event [and therefore an apnea attack] lasts for longer than 45 seconds, the brain might get oxygen deprived and due to this possibility, and alarm system is triggered. A 5V buzzer sounds an alarm, as well as the phone [which is connected to the device via Bluetooth].

### Investigation of Surrounding Factors

This procedure allows for the investigation of the effect of 3 different atmospheric characteristics on the device:

*Temperature* – Leveraging DHT11 Sensor
*Humidity* – Using DHT11 sensor
*CO_2_ Concentration* – Using MQ-135 sensor

Each of the 3 elements being varied are measured in a room, with 3 different subjects suffering from Obstructive Sleep Apnea over a course of 7 days. The data is recorded on to an SD card and analyzed later. The results of the experiment are documented in a later section of this report.

### Android Application

A rudimentary android application was designed that returns the hours of sleep, number of apnea attacks, and Apnea-Hypopnea Index. In addition, it allows for the comparison of AHI on different days under different environmental conditions to assess the effect of said conditions on severity of Apnea, and also sounds an alarm when necessary

## IV RESULTS

3 test subjects known to be suffering from obstructive sleep apnea were taken for testing purposes. The device successfully detected apnea attacks at night. A few false alarms were detected regarding the unusually long apnea events as the device hung up and needed to reset. As for the effects of atmospheric elements, the results are as follows:

### Temperature

The average AHI for the 3 candidates over 2 days was 27 at 18 ° C and 23 at 22 ° C. That is, as temperature increases, AHI decreases

|  | Day 1 AHI<br>[25°C] | Day 2 AHI<br>[22°C] | Day 3 AHI<br>[18°C] |
| --- | --- | --- | --- |
| Subject 1 | 21 | 23 | 28 |
| Subject 2 | 20 | 21 | 23 |
| Subject 3 | 24 | 25 | 30 |

### CO_2_ Concentration

The average AHI for the 3 candidates over 2 days was 23 at 510 ppm and 25.3 at 560 ppm. That is, as [CO2] increases, AHI increases.

|  | Day 1 AHI<br>[440 ppm] | Day 6 AHI<br>[510 ppm] | Day 7 AHI<br>[560 ppm] |
| --- | --- | --- | --- |
| Subject 1 | 21 | 23 | 27 |
| Subject 2 | 20 | 21 | 22 |
| Subject 3 | 24 | 25 | 27 |

